# Timing of meristem initiation and maintenance determines the morphology of fern gametophytes

**DOI:** 10.1101/2021.06.14.448200

**Authors:** Xiao Wu, An Yan, Scott A.M. McAdam, Jo Ann Banks, Shaoling Zhang, Yun Zhou

## Abstract

The alternation of generations in land plants occurs between the sporophyte phase and the gametophyte phase. The sporophytes of seed plants develop self-maintained, multicellular meristems, and these meristems determine plant architecture. The gametophytes of seed plants lack meristems and are heterotrophic. In contrast, the gametophytes of seed-free vascular plants, including ferns, are autotrophic and free-living, developing meristems to sustain their independent growth and proliferation. Compared to meristems in the sporophytes of seed plants, the cellular mechanisms underlying meristem development in fern gametophytes remain largely unknown. Here, using confocal time-lapse live imaging and computational segmentation and quantification, we determined different patterns of cell divisions associated with the initiation and proliferation of two distinct types of meristems in fern gametophytes. Our results reveal how the simple timing of a switch between two meristems has considerable consequences for the divergent gametophyte morphologies of two closely related ferns from Pteridaceae (*Pteris* and *Ceratopteris*). Our result provides evolutionary insight into the function and regulation of gametophyte meristems in seed-free vascular plants.

**Highlight:** Live-imaging of cell growth and division in apical initials and lateral meristems reveals that the timing of a switch between the two meristem identities drives morphology variation in fern gametophytes.

## Introduction

Meristems in land plants are crucial for the formation of the plant body. In seed plants, sporophytes develop multi-cell layered shoot apical meristems (SAMs) and root apical meristems (RAMs) as sustainable resources for the development of shoots and roots, respectively. The reduced gametophytes of seed plants lack meristems and their development is dependent on sporophyte tissues. In seed-free vascular plants, including ferns (Philipson, 1990), gametophytes and sporophytes are independent of each other and gametophytes develop meristems (also called initials) that drive growth and proliferation (Nayar and Kaur, 1971; Imaichi, 2013). Meristems in fern gametophytes persist in an undifferentiated status, self-renew through continuous cell division, produce daughter cells with the potential to differentiate into either photosynthetic cells or cells forming sexual organs (egg-forming archegonia or sperm-forming antheridia), and can eventually terminate in response to environmental signals and developmental cues (Nayar and Kaur, 1971; Banks, 1999; Imaichi, 2013). All of these features suggest that meristems in fern gametophytes share similar functions with the stem cell niches of the sporophytes of seed plants. However, compared to the knowledge on cellular and molecular mechanisms underlying meristem development in seed plants, especially in the model species Arabidopsis (Meyerowitz, 1997; Greb and Lohmann, 2016; Zhou *et al*., 2015; Zhou *et al*., 2018; Han *et al*., 2020), very little is known in ferns. Yet, increasing our understanding of meristems in fern gametophytes could provide important insights in meristem evolution in land plants (Plackett *et al*., 2015, 2018).

There are at least two different types of meristems in the gametophytes of ferns, especially in the species-rich Pteridaceae family (Nayar and Kaur, 1971; Banks, 1999; Conway and Di Stilio, 2020). They consist of the apical initials (with nearest derivatives) and the lateral meristems (that are also called marginal meristems or multicellular meristems) (Nayar and Kaur, 1971; Banks, 1999; Conway and Di Stilio, 2020) (Supplementary Fig. S1). Two derived leptosporangiate ferns, *Ceratopteris richardii* and *Pteris vittata* belong to the Pteridaceae family (PPG, 2016). *Ceratopteris richardii* has been widely used in the studies of physiology, evolution and developmental biology for over 30 years (Hickok *et al*., 1987, 1995; Cooke *et al*., 1995; Eberle *et al*., 1995; Banks, 1999; Chatterjee and Roux, 2000; Plackett *et al*., 2015; Rensing, 2017; Marchant *et al*., 2019; McAdam *et al*., 2016; Geng *et al*., 2021). *Pteris vittata*—the ‘ladder brake’ is unique in its ability to tolerate and hyperaccumulate arsenic, a toxic metalloid element, with exceptionally high efficiency (Ma *et al*., 2001; Dhankher *et al*., 2002; Gumaelius *et al*., 2004; Cai *et al*., 2019). Thus, *P. vittata* provides an important system for studying the transport and metabolism of heavy metals in both sporophytes and gametophytes, and for the phytoremediation of arsenic contaminated soil (Dhankher *et al*., 2002; Gumaelius *et al*., 2004; Ellis *et al*., 2006).

In *C. richardii*, a hermaphroditic gametophyte is composed of a single layer of cells, which eventually develop archegonia and antheridia for sexual reproduction (Banks *et al*., 1993). Several key steps involving the initiation and proliferation of lateral meristems in the hermaphroditic gametophytes of *C. richardii* were characterized using scanning electron microscopy (Banks *et al*., 1993). Through cell growth and developmental analyses, Bartz and Gola (2018) found that the presence of apical initials in *C. richardii* is transient, with these apical initials quickly disappearing after initiation. They also observed and summarized the pattern of prothallus proliferation during the formation of a lateral meristem notch in *C. richardii* (Bartz and Gola, 2018). The evolutionarily conserved meristem regulators in land plants—the LFY family transcription factors, regulate meristem development in *C. richardii* gametophytes and sporophytes (Plackett *et al*., 2018). Recently, an independent study provided a comprehensive view of *C. richardii* gametophyte development that defined distinct stages during the initiation, maintenance and termination of these two types of meristems in this species (Conway and Di Stilio, 2020). These works together demonstrate that hermaphroditic gametophytes of *C. richardii* belong to the classic cordate (heart-shaped) type defined by Nayar and Kaur, 1971, which is caused by the sustained activities of lateral meristems (Banks, 1999; Bartz and Gola, 2018; Conway and Di Stilio, 2020). Different from hermaphrodites, males of *C. richardii* do not have any meristem and they lack active prothallus proliferation, in the presence of antheridiogen—the male-inducing pheromone (Banks *et al*., 1993; Banks, 1999). In contrast, like *C. richardii* hermaphrodites, the prothalli (gametophytes) of species from the genus *Pteris* are also uniseriate, autotrophic and free-living, and generally, they belong to the same cordate type (Nayar and Kaur, 1971; Cai *et al*., 2019). However, species in the genus *Pteris* also develop variable prothallus morphologies that differ from the classic heart or cordate shape (Singh and Khare, 2020), including the model species *P. vittata*.

To date, the potential diversity in gametophyte development and in the patterns of cell divisions within ferns is not known. Any potentially shared or divergent mechanisms in control of apical initials and lateral meristems in species from this family are yet to be identified. Furthermore, there has been no developmental explanation for how variable prothallus morphologies are present in ferns. To tackle these questions, we took both snapshots and time-lapse live images of developing gametophytes of *P. vittata* using confocal microscopy, performed computational segmentation and image analysis, and quantitively examined the patterns of cell division and growth in *P. vittata* with the comparison to *C. richardii*. Our results reveal unique patterns of cell division that are closely associated with the initiation and proliferation of apical initials and lateral meristems. The variable timing of the maintenance of apical initials and lateral meristems determines the variable gametophyte morphology of *P. vittata*, which is different from the classical cordate shape of *C. richardii*. Our work provides new insights into the function and regulation of meristems in fern gametophytes, and it provides a new quantitative platform for future studies into the cellular mechanisms underlying diverse gametophyte morphologies more broadly across ferns (Christenhusz and Byng, 2016).

## Materials and Methods

### Plant materials and growth condition

The spores of *C. richardii* (Hn-n) and *Pteris vittata* were described previously in (Banks *et al*., 1993; Cai *et al*., 2019), respectively. The spores were sterilized with the 33% bleach and 0.5% Tween20 and rinsed five times with sterile water, then kept in dark for two days, as described previously (Banks *et al*., 1993; Cai *et al*., 2019). After that, spores were spread on the FM medium that contains 0.5x MS salts (Phytotech), pH 6.0, 0.7% agarose (Sigma). The gametophytes of *C. richardii* and *Pteris* were grown in the Percival growth chambers at 28°C with continuous light and 80% humidity.

### Confocal live imaging and image analysis

The gametophytes of *C. richardii* and *Pteris* were live imaged using a Zeiss LSM 880 upright confocal microscope. In the presence of antheridiogen, the male-inducing pheromone, male gametophytes of *C. richardii* do not develop any lateral meristem (Banks et al., 1993; Banks, 1999). In addition, *C. richardii* males just transiently maintain the wedge-shaped apical initial (Banks, 1999; Bartz and Gola, 2018; Conway and Di Stilio, 2020). For these reasons, only hermaphroditic gametophytes of *C. richardii* were imaged (shown in Figure 1) and *C. richardii* males were not included in this study. In the current growth conditions and within the time frame examined (3 to 32 days after inoculation, DAI), all the imaged *P. vittata* gametophytes did not produce gametangia, nor were they sexually dimorphic.

**Fig. 1.**
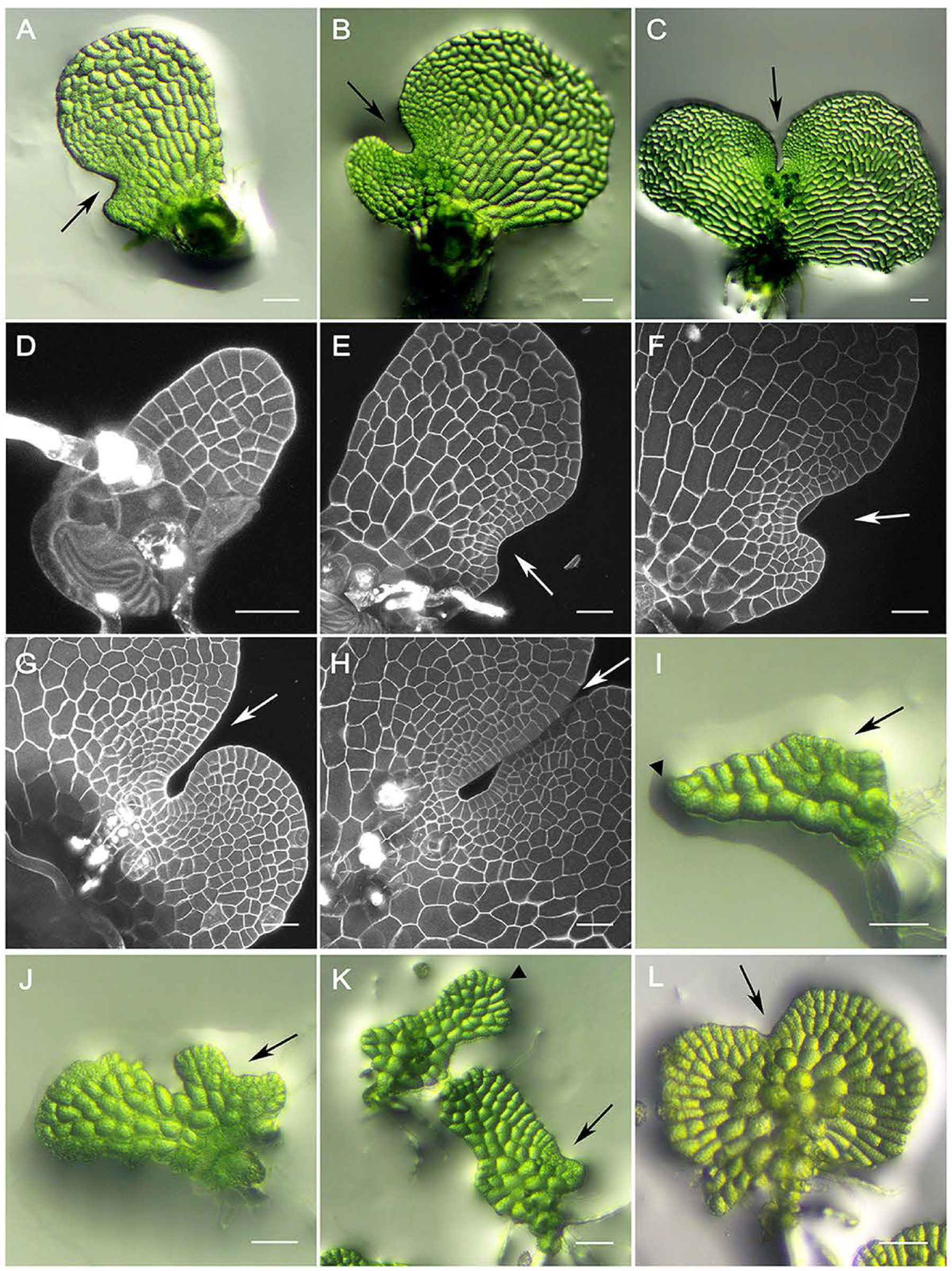
The prothalli of *Ceratopteris richardii* (A-H) and *Pteris vittata* (I-L). (A-C) Representative images of hermaphroditic gametophytes of *C. richardii* at 8 (A). 10 (B) and 12 (C) days after inoculation (DAI). (D-H) Confocal imaging of representative hermaphroditic gametophytes of *C. richardii* at 5 (D), 7 (E), 8 (F), 10 (G) and 12 (H) DAI. (I-L) Representative images of *P. vittata* gametophytes at15 DAI (I-K) and 26 DAI (L). Triangle arrow indicates the pointed growing tip. Arrow indicates the lateral meristem notch. Within the time frame examined, *P. vittata* gametophytes do not develop gametangia. (I) represents the Type I, (J) and the lower sample in (K) represent the Type II, and the upper sample in (K) represents the Type III summarized in Table 2. (L) represents the Cordate, Ceratopteris-like gametophyte summarized in Table 1. Scale bar: 10 µm (A-C, I-L), 50 µm (D-H).

To quantitively determine the initiation and development of apical initials and lateral meristems, different *Pteris vittata* gametophytes were imaged every day as snapshots. The number of samples imaged was listed in Supplementary Table S1. The confocal snapshots of the samples from 3 DAI to 32 DAI were included in this study. After 32 DAI, Pteris gametophytes curve from the wings of prothalli, making it technically challenging and inaccurate to perform confocal imaging and 2D computational analysis.

To determine the dynamic cell growth and division in apical initials and lateral meristems, Pteris gametophytes were live imaged over the 12-hour or 24-hour period. The non-invasive time-lapse live imaging was performed on the gametophytes grown on the FM plates. Before taking the confocal imaging, the FM plates with the gametophytes were moved out of the growth chamber, and the *P. vittata* gametophytes were stained with propidium iodide (PI) and rinsed with sterilized water twice. After taking the first time point, the gametophytes were transferred to freshly made FM plates, and then moved back to the growth chamber and cultured in the same condition until the second time point.

The settings of confocal imaging were described previously in detail (Geng and Zhou, 2019) with minor modifications in this study. Specifically, in the time-lapse experiment, PI was excited by a 561 nm laser line. Other specific parameters in the Zeiss ZEN software were listed, including the Scan mode: frame; Frame size: 512 × 512; Scanning speed: 8-10; Scanning direction: bi-direction; Averaging number: two; Averaging method: mean; Bit depth: 16 Bit; and Scanning Interval: 1 µM. The confocal images were processed using the Fiji/ Image J software to generate the z projection view.

The 2D image segmentation of the confocal images was performed using the 2D watershed method as described previously (Vincent and Soille, 1991). In all the confocal images of this study, the cell wall signal is much brighter than the cytosolic signal (Supplementary Fig. S3A), which fits the previously published watershed algorithm (Vincent and Soille, 1991). The 2D watershed was carried out using a MATLAB software built-in implementation (MATLAB 2019a). The code is available upon request. As the result of 2D watershed segmentation, each segmented cell was designated a unique label as shown in Supplementary Fig. S3B and Supplementary Fig. S11. To quantify the area of each segmented cell, the total pixel numbers were first summed for each area with unique label and then multiplied with the pixel area size, which were also implemented in the MATLAB software. The areas of each segmented cell from individual images were then exported as individual tables (as shown in Supplementary Tables S2-S8). For the visualization, the size of each segmented cell was quantitatively indicated by color with the range specified in each figure legend.

## Results

### The morphology of the Pteris vittata prothallus is highly variable compared to that of Ceratopteris richardii

As described previously (Banks, 1999; Conway and Di Stilio, 2020), *C. richardii* hermaphroditic gametophytes are cordate (Fig. 1A-H), like the majority of terrestrial fern gametophytes (Nayar and Kaur, 1971). In *C. richardii*, the apical initial of a germinating gametophyte disappears quickly and before the establishment of the lateral meristem (Fig. 1A, D-E) (Bartz and Gola, 2018). Once initiated, the lateral meristem of *C. richardii* actively proliferates, forming the meristem notch that first appears as a shallow concave at 7 days after inoculation (DAI) and then leads to a heart-shape structure at 12 DAI (Fig. 1E-H). *C. richardii* gametophyte development is stereotypical in that there is little variation in the development and the overall size and shape of mature *C. richardii* prothalli is stable (Fig. 1A-H) (Banks *et al*.,1993; Banks, 1999; Conway and Di Stilio, 2020). Unlike *C. richardii, Pteris* gametophyte development is variable (Fig. 1I-L, Supplementary Fig. S2). Mature *P. vittata* prothalli are often cordate (Fig. 1L) (Cai *et al*., 2019), although more than one third of gametophytes examined have a non-cordate shape at 26 DAI (Table 1). In addition, before reaching the maturity, *P. vittata* prothalli display highly variable shapes (Fig. 1I-L; Supplementary Fig. S2). Some prothalli form a rectangular plate (18 out of 51, Table 2) (Fig. 1K), some develop a rectangular plate associated to a shallow notch or a small heart-shape structure (Fig. 1J, K) (20 out of 51, Table 2), while some maintain a pointed tip and a meristem notch at the same time (Fig. 1I) (13 out of 51, Table 2). Since the meristems drive gametophyte growth and development in ferns, these observations (Fig. 1, Tables 1, 2) led us to explore whether the distinctive gametophyte morphologies between *C. richardii* and *P. vittata* are due to the differential activity and/or regulation of meristems.

**Table 1.** Variable morphology of the *P. vittata* gametophytes at 26 DAI.

| Total gametophytes observed at 26 DAI | Non-cordate, different from Ceratopteris | Cordate, Ceratopteris-like | % of gametophytes different from Ceratopteris | % of gametophytes, Ceratopteris-like |
| --- | --- | --- | --- | --- |
| 40 | 15 | 25 | 37.50% | 62.50% |

**Table 2.** Variable morphology of the *P. vittata* gametophytes at 15 DAI.

| Total gametophytes observed at 15 DAI | Type I | Type II | Type III |
| --- | --- | --- | --- |
| 51 | 13 | 20 | 18 |
| Total | % of Type I | % of Type II | % of Type III |
| 100% | 25.49% | 39.22% | 35.29% |

Note.
Type I: prothallus with a pointed tip;
Type II: prothallus with a rectangular plate and a notch;
Type III: prothallus with a rectangular plate or a rectangular plate-like structure.

### Snapshots and computational analysis reveal the variable apical initials and invariable lateral meristems in P. vittata

To capture the variability across *P. vittata* gametophytes, we first imaged prothalli at different developmental stages and quantitatively defined the initiation, maintenance and termination of the two types of meristems (the apical initial and the lateral meristem) (Fig. 2A-X, Supplementary Figs. S3-S11). *P. vittata* spores were plated on solidified growth medium, and different individual gametophytes were imaged every day using laser scanning confocal microscopy (Supplementary Table S1), from spore germination (three days after inoculation, DAI) to a late developmental stage with the fully expanded blade (32 DAI) (See methods for details). A computational analysis of each confocal image was performed to define each cell and quantify cell area in each gametophyte selected (Supplementary Figs. S3, S11, Supplementary Tables S2-S8).

**Fig. 2.**
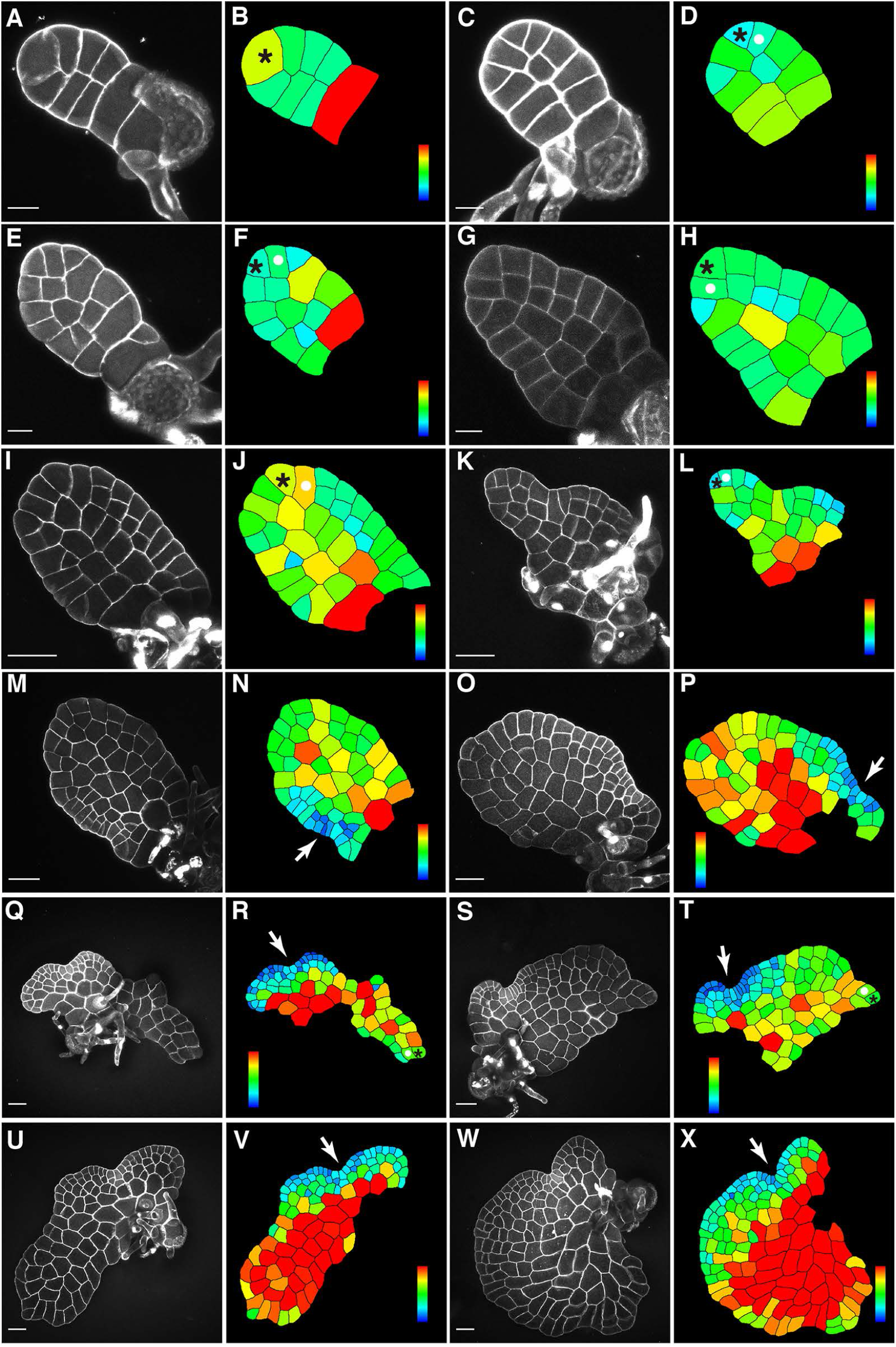
The confocal imaging and computational analysis of *P*.*vittata* gametophytes at different developmental stages. Twelve representative garnetophytes of *P. vittata* were stained and imaged respectively at 6 DAI (A), 7 DAI (C), 8 DAI (E), 9 DAI (G), 11 DAI (I), 15 DAI (K), 17 DAI (M), 19 DAI (0), 22 DAI (Q and S), 27 DAI (U) and 31 DAI (W) through laser scanning confocal microscopy. (B, D, F, H, J, L, N, P, R, T, V, X) show the computational segmentation and cell size quantification of confocal images in (A, C, E, G, I, K, M, O, Q, S, U, W). Stars indicate the apical initials, white dots indicate the trapezoid-shaped cells, and arrows indicate the lateral meristems. Scale bar: 20 µm in (A, C, E, G) and 50 µm in (I, K, M, O, Q, S, U, W). Gray (A, C, E, G, I, K, M, O, Q, S, U, W): propidium iodide (PI) stain. Color bar indicates the quantified area of each segmented cell with the scale from blue (0) to red (at or above 1000 µm^2^) in (B, D, F, H, J, L), with the scale from blue (0) to red (at or above 1300 µm^2^) in (N, P), and with the scale from blue (0) to red (at or above 1600 µm^2^) in (R, T, V, X). At least three independent biological replicates showed the pattern similar to each snapshot included in the figure. Within the time frame examined,*P. vittata* garnetophytes do not develop gametangia. The snapshots captured the different stages of apical inilial development, including its initiation (A-8), proliferation (C-L, Q-T) and termination (M-P, W-X). (K-L) represent the Type I summarized in Table 2. Additional confocal images and associated quantification analyses are included in Supplementary Figs. S4-S9.

Since 3 DAI, *P. vittata* undergoes the spore germination process as described (Nayar and Kaur 1971), forming an elongated filament along the apical-basal axis (Supplementary Fig. S4). At as early as 6 DAI, the wedge-shaped initial cell (Fig. 2A-B, Supplementary Fig. S5A-B, marked by star) at the apical tip is morphologically distinguishable from all the other cells of the prothallus (Supplementary Table S1). Interestingly, we found that the timing in maintaining the apical initials is highly variable: in a number of gametophytes at different DAI, the morphological feature of apical initials is maintained at the growing tip, showing the pattern of one wedge-shaped cell outside and one trapezoid-shaped cell inside (Fig. 2C-L, Q-T, Supplementary Figs S5C-H, S6A-P); in contrast, in many other gametophytes, the morphological feature of apical initials disappears, showing a pattern of one rectangle-shaped cell outside and one wedge-shaped cell inside (Fig. 2O-P, Supplementary Figs. S6W-X, S8I-J, S9M-N), which suggests the termination of apical initials.

Different from apical initials, we found the growth pattern to initiate and maintain a lateral meristem in *P. vittata* is less variable (Fig. 2, Supplementary Figs. S6-S10). The lateral meristem of *P. vittata* is morphologically visible at 10-11 DAI (Supplementary Fig. S8A-D). At one lateral side, the prothallus forms a cluster of elongated rectangle cells that serve as the initiation site of the lateral meristem notch (Fig. 2M-P, Supplementary Fig. S8A-J). Importantly, the results from the computational segmentation and quantification demonstrated that the small size (quantitatively indicated by various shades of blue color) is the conserved feature for these cells of lateral meristems (Supplementary Tables S2-S8), which can be found in almost all the gametophytes in the snapshots (Fig. 2, Supplementary Figs. S8, S9). Once initiated, these small cells continuously proliferate, forming the meristem notch from one lateral side of prothallus (Fig. 2M-X, Supplementary Figs. S8A-L, S9C-P), similar to the pattern observed in *C. richardii* (Fig. 1A-D) (Banks *et al*., 1993, Conway and Di Stilio, 2020).

### Co-existence of an apical initials and a lateral meristem during the P. vittata gametophyte development

To comprehensively understand the dynamics of apical initials and lateral meristems during gametophyte development, we examined all the confocal images of different gametophytes and calculated the percentages of gametophytes with either an apical initial or a lateral meristem at indicated DAI (Fig. 3, Supplementary Table S1). In general, the apical initials become morphologically distinguishable in the gametophytes at 5-6 DAI. The percentage of gametophytes that maintain the apical initials increases quickly, reaching a peak of almost 100% at 9-10 DAI. After that, this percentage gradually decreases to ∼ 22% at 31-32 DAI. Compared to the apical initials, the initiation and proliferation of lateral meristems has a different but overlapping trend in developing gametophytes. Lateral meristem cells are morphologically distinguishable from other cells at 9-10 DAI. After that, the percentage of gametophytes with lateral meristems increases steadily, reaches the peak of 100% at 23-24 DAI and remains unchanged (until 31-32 DAI when experiments were completed). These quantitative results demonstrate that a large portion of gametophytes maintain both apical initials and lateral meristems for a long time (Supplementary Table S1), which is distinct from the pattern observed in *C. richardii* (Fig. 1 A-D) (Banks *et al*., 1993, Conway and Di Stilio, 2020). This result suggests that *P. vittata* has more indeterminate growth, and this could explain the differences in gametophyte shapes between cordate *C. richardii* gametophytes and those of *P. vittata*.

**Fig. 3.**
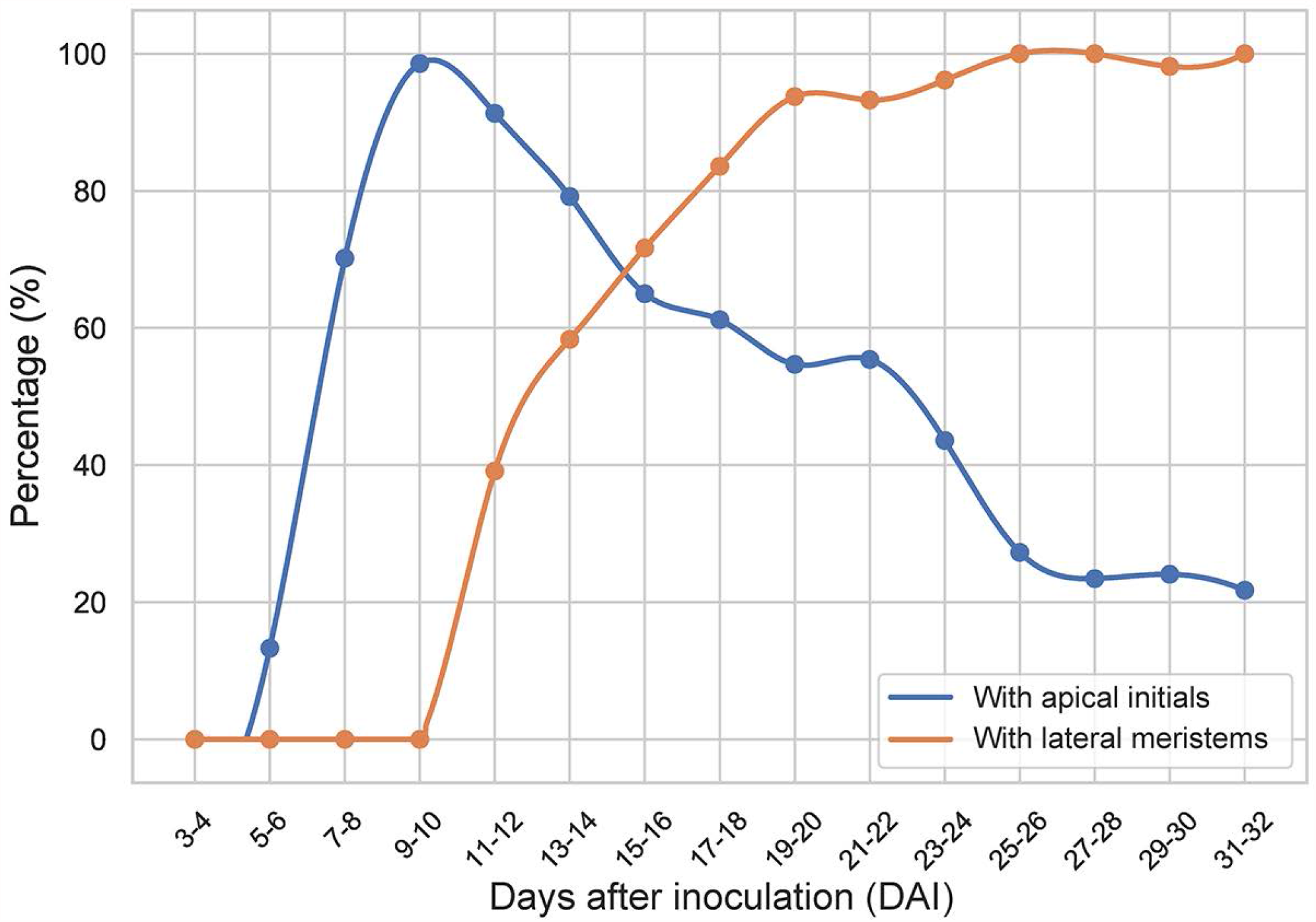
The dynamics of both apical initials and lateral meristems during gametophyte development in *P. vittata*, Blue represents the percentage of gametophytes that maintain the apical initial at the indicated days after inoculation (DAI). The presence (including the stages of initiation and proliferation) of an apical initial is determined by its distinct morphology-showing a wedge-shaped cell at the apical tip of a gametophyte. As indicated in Fig.2A-L and Fig. S5A-H The absence (termination) of an apical initial is also determined based on the loss of morphological characteristics-showing a trapezoid-shaped cell at the apical tip of a gametophyte, like the ones in Fig. 2O-P and Fig. S6W-X. At 3 DAI. the spore coat starts to crack open and only part of spore cell and a developing rhizoid can be observed At 6 DAI, the wedge-shape apical initial can be observed. Red represents the percentage of gametophytes that maintain the lateral meristems at indicated DAI. The source data for this figure are included in Supplementary Table S1.

### Apical initials are maintained in the P. vittata gametophytes with variable non-cordate morphologies

We then found that all the non-cordate gametophytes maintain the wedge-shaped cells (marked by stars) at their apical tips, suggesting an important role of apical initials in shaping *P. vittata* prothalli. For example, in the snapshots, some “pyramid-shaped” gametophytes do not yet form lateral meristems but maintain apical initials at the growing tip along the apical-basal axis (Fig. 2K-L, Supplementary Figs. S7A-B, S9A-B), showing an apical initial dominated growth pattern. Some gametophytes consist of two distinct parts, including one “pyramid-shaped” region with a wedge-shaped apical cell and one heart-shape region with a lateral meristem notch (Fig. 2Q-R, Supplementary Figs. S7G-H, S9C-H). These results suggest that both apical initials and lateral meristems have been active, forming two growing points at different positions of prothalli. At late developmental stages (over 30 DAI), the gametophytes with both apical initials and lateral meristems form the notch (marked by blue color) at one lateral side of the blades but also maintain the pointed growing tips at the apical part (Fig. 2S-T, Supplementary Fig. S9I-L). In contrast, the gametophytes with lateral meristems but lacking apical initials show the heart shape (Fig. 2U-X, Supplementary Fig. S9M-P), comparable to that in *C. richardii* (Fig. 1B, E-G). We interpret that the sustained activity of apical initials in *P. vittata* drives the polarized tip growth of prothalli along the apical-basal axis, resulting in the non-cordate shapes. In addition, the variable timing in the termination of apical initial activity likely contributes to the variable morphologies of prothalli at maturity.

### Oblique cell division during the initiation and proliferation of apical initials in P. vittata gametophytes

To quantitively determine the patterns of cell division in apical initials of *P. vittata* gametophytes, we performed confocal time-lapse live imaging and the computational image analysis (Figs. 4-7). We first imaged the living gametophytes at an early developmental stage (10 DAI) when their apical initials have been established but their lateral meristems are not yet initiated (Fig. 4). The wedge-shaped apical initial located at the apical tip of the prothallus (marked by star) is morphologically distinguishable from all the other cells (Fig. 4A-C, G-I). Once specified, the apical initials are self-maintained through cell division (Fig. 4A-M). Over a 12-hour interval, the new cell wall is formed obliquely in the middle of the wedge-shaped apical initials (Fig. 4A-M). This type of oblique division results in two daughter cells with distinct shapes: a new wedge-shaped apical cell outside and a trapezoid-shaped cell inside (Fig. 4D-F, J-L, M). The newly formed apical cell is divided again by new wall, which is perpendicular (or near perpendicular) to the previous oblique wall (Fig. 4G-L, N). In contrast, the trapezoid-shaped daughter cell (Fig. 4G-L, O) undergoes periclinal division, resulting in two new trapezoid-shaped cells (Fig. 4G-L, O). This dynamic division pattern revealed by the time-lapse imaging and image analysis is consistent with the shapes of the apical initials and their immediate progeny observed in multiple snapshot images (Fig. 2, Supplementary Figs. S5-S10).

**Fig. 4.**
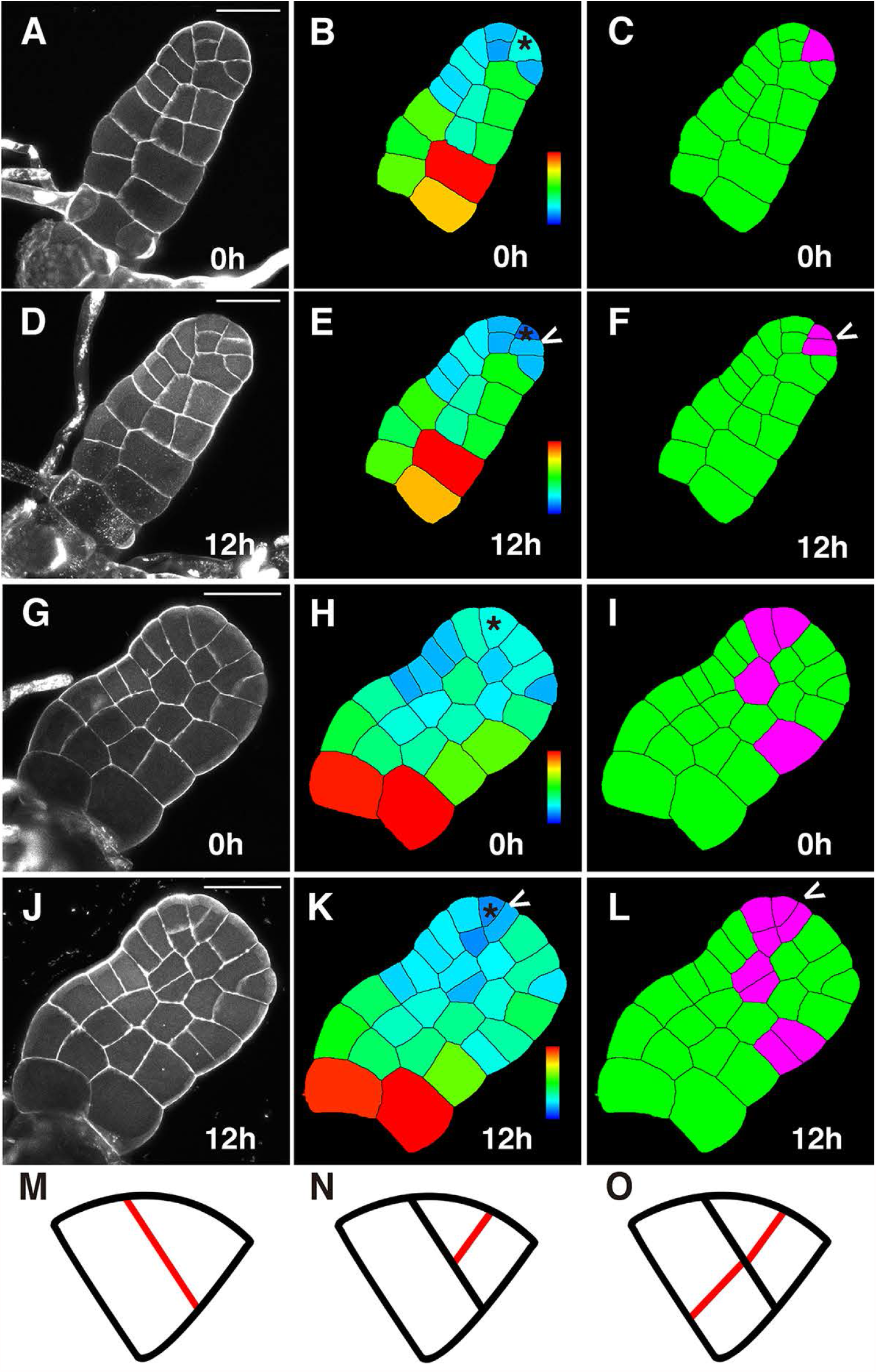
The patterns of cell divisions in the apical initials of *P*.*vittata* gametophytes at 10 DAI. Two gametophytes (A-F,G-L) were stained and live imaged through the laser scanning confocal microscopy at 0 hour (A, G) and 12 hours (D, J). (B,E, H, K) show the computational segmentation and cell size quantification of confocal images in (A, D,G,J), respectively. (C,F, I, L) highlight cell divisions in the time-lapse images in (A, D,G,J) over 12 hours, with dividing cells in purple and non-dividing cells in green. Star indicates the wedge-shaped apical initial and ‘V’indicates cell division in the apical initials. (M-O) Diagrams and illustration of cell division patterns in the apical initial and its derivatives, with newly formed cell wall in red. Scale bar (A, D,G,J): 50 µm. Gray (A,D, G, J): propidium iodide (PI) stain. Color bar in (B, E, H,K) indicates thequantified area of each segmented cell with the scale from blue (0) to red (at or above 1600 µm^2^). 15 independent biological replicates (10 DAI) in the time-lapse experirnent showed the similar patterns of cell division in apical initials.

### Anticlinal cell division during the initiation of lateral meristems in P. vittata gametophytes

We also performed the time-lapse live imaging to determine the specific pattern of cell division and growth that potentially initiates the lateral meristems (Fig. 5A-O). Over 12 hours, we were able to capture the earliest event during the formation of a lateral meristem, with the anticlinal cell division simultaneously occurring in multiple adjacent cells on one lateral side of each prothallus (Fig. 5A-L, N, O). Through the computational segmentation of the confocal images and the subsequent quantification of cell area (Fig. 5B-C, E-F, H-I, K-L), we found that this unique division pattern leads to a doubling of cell number and a cluster of adjacent rectangular cells with small size, which are quantitatively indicated by the various shades of blue color. These blue-colored cells (Fig. 5E, K) serve as the initiation site of lateral meristem development that we have repeatedly observed in a number of *P. vittata* gametophytes in the snapshot images (Fig. 2N, P, Supplementary Fig. S8B, D, H, J). Besides this unique pattern of anticlinal division in multiple adjacent cells at one side of the blade, we also noticed periclinal cell divisions on the other lateral side of the blade (Fig. 5A-L, M) and random cell divisions at the center of the prothallus (Fig. 5A-F, M), both of which seem to be uncoupled from the lateral meristem initiation, but contributing to the expansion of prothalli. All these results suggest that specific patterns of cell division are related to the cell fate specification in fern gametophytes, and at early stages of *P. vittata* gametophyte development, cells outside of the meristem initiation site also maintain division activity.

**Fig. 5.**
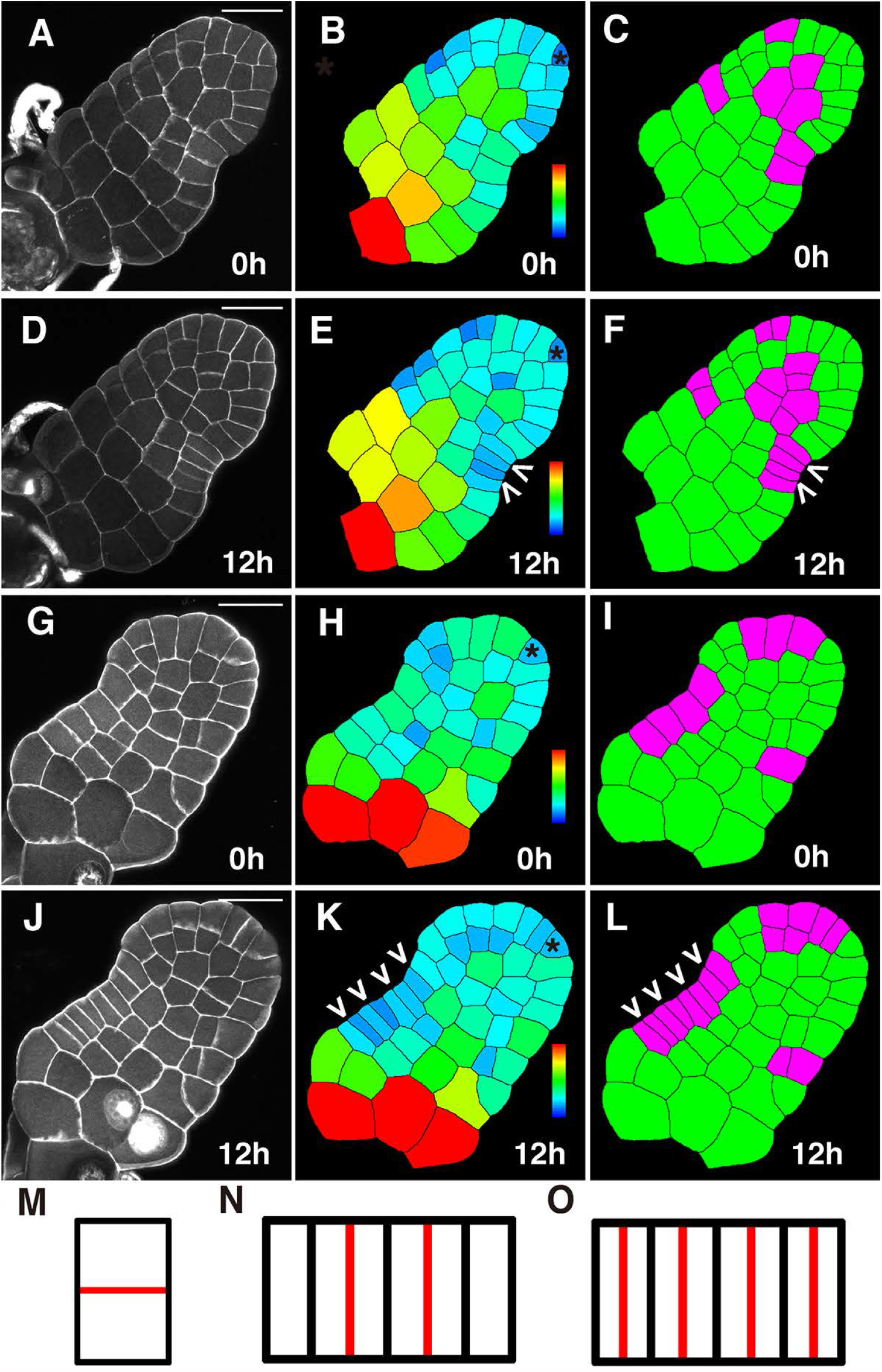
The paterns of cell divisions during the initiation of lateral meristems in *P. vittata* gametophytes at 10 DAI. Two gametophytes (A-F, G-L) were stained and live imaged through the laser scanning confocal microscopy at 0 hour (A, G) and 12 hours (D, J). (B, E, H, K) show the computational segmentation and cell size quantification of confocal images in (A, D, G, J), respectively. (C, F, I, L) highlight cell divisions in the time-lapse images in (A, D, G, J) over 12 hours, with dividing cells in purple and non-dividing cells in green. Star indicates the wedge-shaped apical initial and ‘V’ indicates cell division during the initiation of lateral meristems. (M-O) Diagrams summarize the periclinal (M) and anticlinal cell divisions (N-O) during the lateral meristem initiation, with newly formed cell wall in red. Scale bar: 50 µm. Gray (A, D, G, J): propidium iodide (PI) stain. Color bar in (B, E, H, K) indicates the quantified area of each segmented cell with the scale from blue (0) to red (at or above 1600 µm^2^). Six independent biological replicates (10 DAI) in the time-lapse experiment showed the similar patterns of cell divisions during initiation of lateral meristems.

### Patterns of cell division during the proliferation of lateral meristems

To further determine the specific pattern of cell division and growth that potentially drives the lateral meristem proliferation, we live imaged the *P. vittata* gametophytes at a later developmental stage (27 DAI) (Fig. 6), with the established meristem notch. Over a 24-hour period, anticlinal cell divisions actively occur in multiple cells at the lateral meristem notch, leading to the increase of cell number in the outmost layer and the proliferation of lateral meristems (Fig. 6A-H). More importantly, through image analysis, we found the reverse “T” type of cell division pattern (described in Imaichi, 2013) that is closely associated with the cells at the center of lateral meristems (Fig. 6A-M). Specifically, this pattern consists of the periclinal cell division followed by the anticlinal cell division, resulting in three daughter cells from one initial (Fig. 6A-M). In addition, we found a number of cells, most of which located at outermost layers, undergo two or multiple rounds of cell division over 24 hours in *P. vittata* gametophytes (Fig. 6), whereas only one round of cell division was observed within the 12-hour period (Fig. 4, 5).

**Fig. 6.**
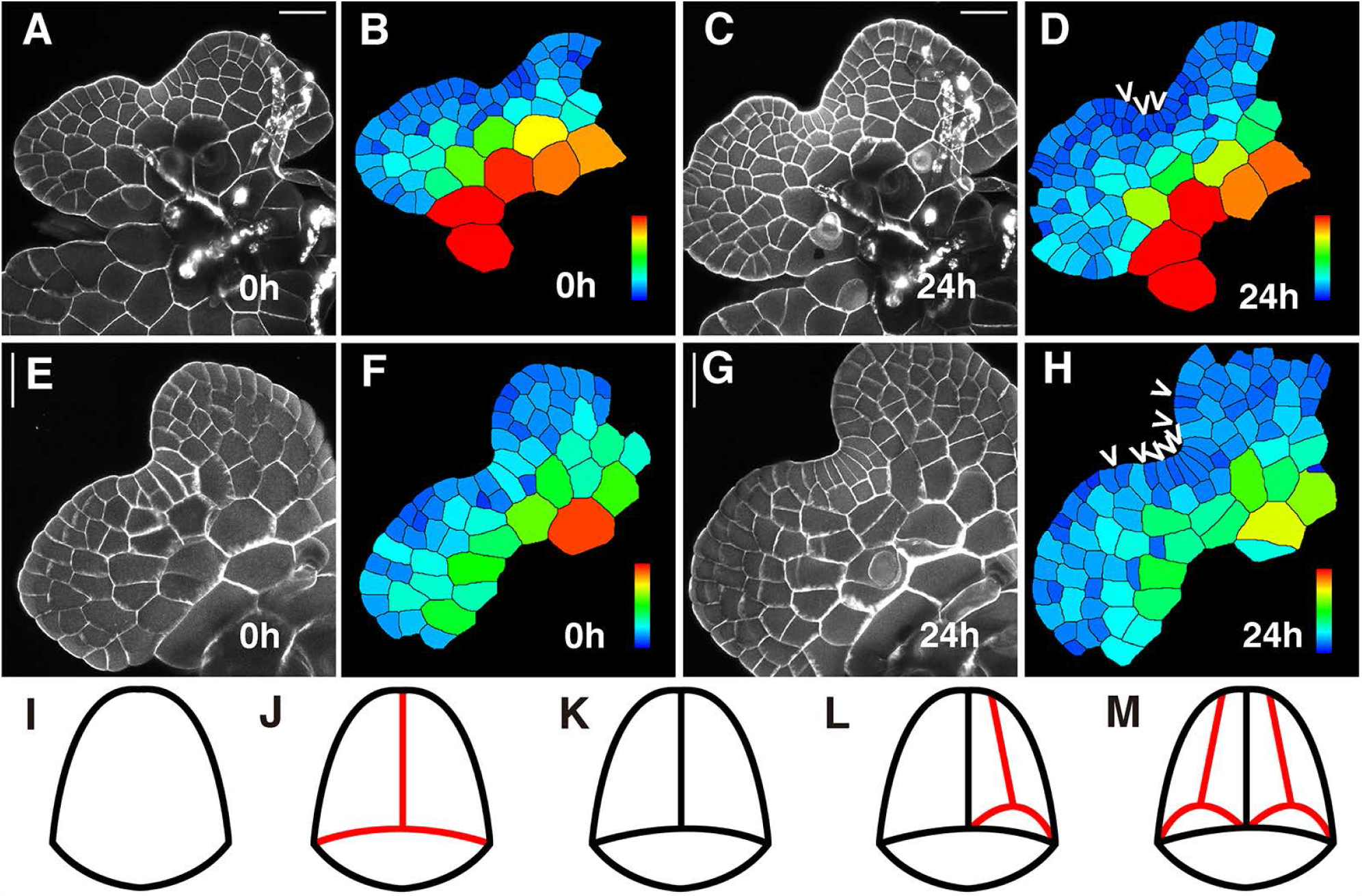
The patterns of cell divisions during the proliferation of lateral meristems in *P. vittata* gametophytes at 27 DAI. Two gametophytes (A-D, E-H) were stained and live imaged through the laser scanning confocal microscopy at 0 hour (A, E) and 24 hours (C, G). (B, D, F, H) show the computational segmentation and cell size quantification of confocal images in (A, C, E, G), respectively. ‘V’ indicates cell division in the notch of lateral meristems. (I-M) Diagrams summarize the patterns of cell divisions in the lateral meristems, with newly formed cell wall in red. Scale bar. 50 µm. Gray (A, C, E, G): propidium iodide (PI) stain. Color bar indicates the quantified area of each segmented cell with the scale from blue (0) to red (at or above 2400µm^2^). 18 independent biological replicates (27 DAI) in the time-lapse experiment show the similar patterns of cell divisions during the proliferation of lateral meristems.

### Cell divisions simultaneously occur in both apical initials and lateral meristems

As we described in Fig. 3, at late developmental stages (27 DAI, for example), a portion of *P. vittata* gametophytes still maintain both lateral meristems and apical initials (Fig. 7). We then performed the time-lapse live imaging to directly view and quantitatively analyze the cell division patterns in these *P. vittata* gametophytes with variable morphologies. Over a 24-hour period, the oblique cell division occurs in the apical initials, leading to the continuous proliferation of the wedge-shaped initial cells and their derivatives at the tip of gametophytes. At the same time, the reverse “T” pattern of cell division occurs in the lateral meristems, driving the proliferation of the meristem notch at the side of the blade. These results not only show the maintenance of an apical initial and a lateral meristem during gametophyte development, but also demonstrate the active roles of both meristems in shaping the gametophytes.

**Fig. 7.**
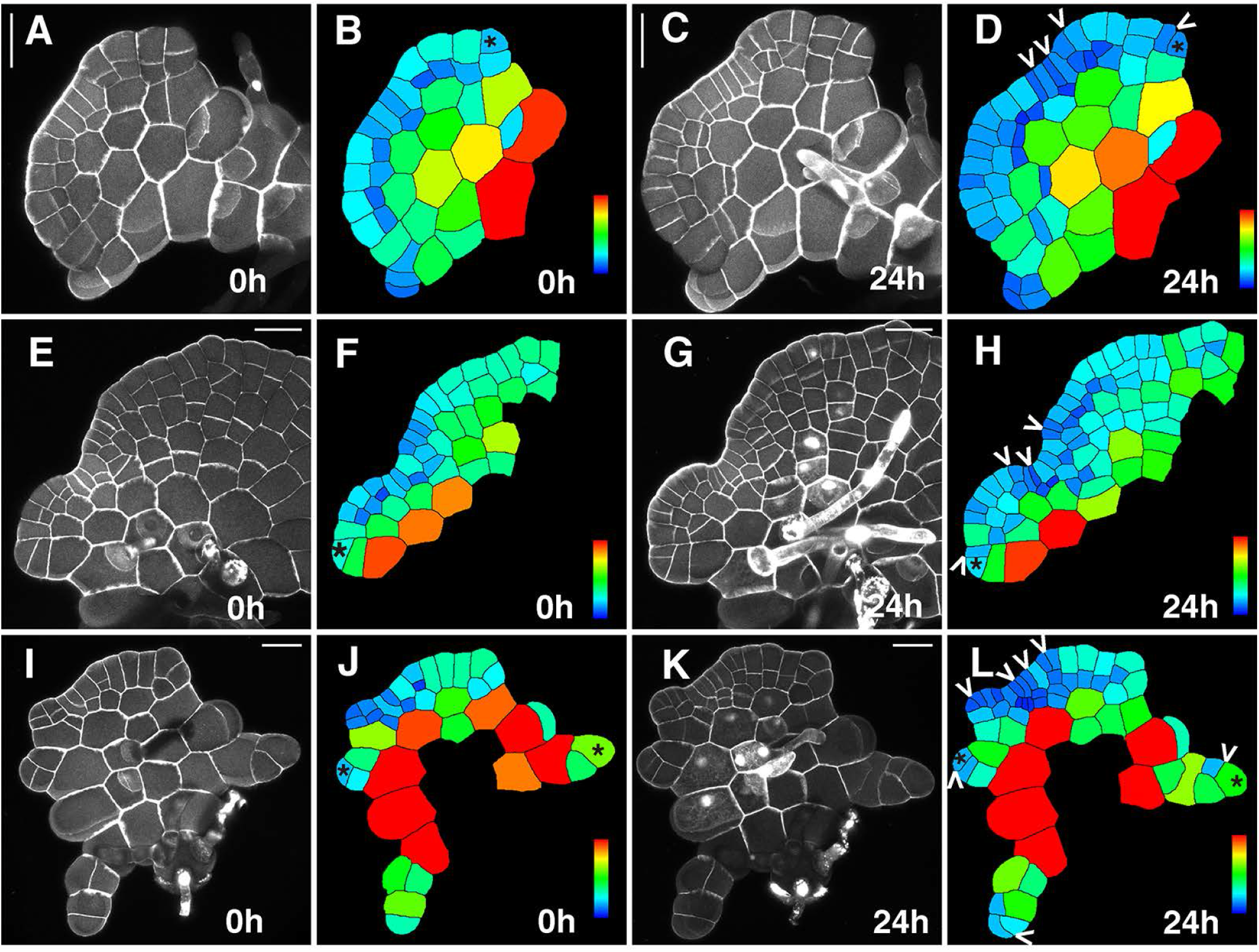
Cell divisions in both apical initials and lateral meristems of *P. vittata* gametophytes at 27 DAI. Three gametophytes (A-D, E-H, I-L) were stained and live imaged through the laser scanning confocal microscopy at 0 hour (A, E, I) and 24 hours (C, G, K). (B, D, F, H, J, L) show the computational segmentation and cell size quantification of confocal images in (A, C, E, G, I, K), respectively. Star indicates the apical initial and ‘V’ indicates cell division in the apical initials or in the notch of lateral meristems. Scale bar: 50 µm. Gray (A, C, E, G, I, K): propidium iodide (PI) stain. Color bar indicates the quantified area of each segmented cell with the scale from blue (0) to red (at or above 1600 µm^2^) in (F, H) and with the scale from blue (0) to red (at or above 2400 µm^2^) in (B, D, J, L). Five independent biological replicates (27 DAI) in the time-lapse experiment show the similar patterns of cell divisions in both apical initials and lateral meristems.

## Discussion

As sister to seed plants, ferns represent an important node in the phylogeny of land plants (Plackett *et al*., 2015). Ferns share derived traits with seed plants, whereas they produce dormant spores instead of seeds and they develop gametophytes that are independent of sporophytes, the ancestral traits shared with other non-seed plant lineages. Therefore, ferns are key to understanding the evolution of plant body formation (Plackett *et al*., 2015). This study using two related and important ferns provides crucial knowledge of meristem development in gametophytes. Future studies into the broad diversity of gametophyte morphologies across the fern clade (Christenhusz and Byng, 2016), using the approach we have established here, will provide the more comprehensive view on meristem function and fern evolution.

### Cell division patterns and dynamics of initial cells

Through confocal time-lapse live imaging, computational segmentation, and image analysis, we determine the distinct patterns of cell divisions in the apical initial and in the lateral meristem. The oblique cell division in an apical initial gives rise to one wedge-shaped daughter cell outside and one trapezoid-shaped daughter cell inside (Figs. 4, 7). This type of cell division is associated with the initiation and self-maintenance of apical initials. In contrast, seen from multiple snapshot images, when the gametophytes develop one trapezoid-shaped cell outside and one wedge-shaped cell inside at the apical side (Fig. 2O-P, Supplementary Figs. S6W-X, S8I-J, S9M-N), the morphological signature of apical initials disappears. This pattern is likely resulted from the periclinal cell division in the apical initial, associated with the termination of this type of meristems.

Unlike apical initials, the simultaneous anticlinal cell division in adjacent cells at one lateral side of gametophytes is a unique feature that defines the initiation of lateral meristems (Fig. 5A-L, N-O). In addition, the typical reverse “T” pattern (with a periclinal division followed by an anticlinal division) drives the maintenance of initials in lateral meristems (Fig. 6A-M). Taken together, unique patterns of cell divisions in gametophytes directly associate with different cell identities, suggesting that cell division likely plays a role in control of meristem behaviors in fern gametophytes. This finding in fern gametophytes is different from the knowledge gained in *Arabidopsis* shoot apical meristems (Shapiro *et al*., 2015), because patterns of cell division in *Arabidopsis* SAMs do not contribute to the cell fate specification. Future work to uncover the molecular basis of these division patterns in ferns will provide more understanding on the initiation and maintenance of different types of gametophyte meristems.

### Lateral meristems and apical initials together define the morphology of gametophytes in P. vittata

In *C. richardii*, apical initials are just transiently maintained and quickly disappeared in hermaphroditic gametophytes (Banks, 1999; Bartz and Gola, 2018). The lateral meristems in *C. richardii* hermaphrodites play a major role in proliferating prothalli and give their heart shape appearance at maturity (Banks *et al*., 1993, Banks, 1999; Conway and Di Stilio, 2020). In contrast, the apical initials and lateral meristems together determine the morphology of *P. vittata* gametophytes at maturity. Through taking the snapshots of ∼900 gametophytes at different DAI (Fig. 3, Supplementary Table S1), we found that the majority of *P. vittata* gametophytes contain both apical initials and lateral meristems at the same time (Figs. 2, 3, Supplementary Figs. S5-S9) and the apical initials in *P. vittata* gametophyte can be maintained for a much longer time through the continuous oblique cell division (Figs. 4, 7). The apical initial dictates polarized growth along the apical-basal axis, whereas the lateral meristem activity drives the formation of a meristem notch from one side of gametophytes and promotes lateral growth of blades (Figs. 4-7). The co-existence of these two types of meristems promote the growth of prothalli in two different directions, therefore leading to variable shapes and morphologies of gametophytes (Fig. 2, Supplementary Figs. S7, S9). The timing of a switch from the activity of apical initials to the function of lateral meristems is different between *C. richardii* and *P. vittata*, determining whether a gametophyte eventually develops a cordate or non-cordate shape. While cordate gametophytes are common in terrestrial ferns, many other fern species show highly variable morphology of gametophytes, ranging from the classic heart-shape to indeterminately growing strips or ribbon-like gametophytes (Nayar and Kaur, 1971; Imaichi, 2013). A variable gametophyte morphology often results in a very long-lived gametophyte stage, exemplified in the extreme by the number of fern species that are found as only gametophytes, with highly modified filamentous morphology (Pinson *et al*., 2017). In addition, the variation in gametophyte morphology has been associated with differences in ecological tolerance and strategies, with the indeterminately growing gametophytes of a number of lineages associated with desiccation tolerance and a capacity to reproduce in epiphytic niches (Watkins and Cardelús, 2012; Watkins *et al*., 2007). It will be interesting to explore whether the simple changes of meristem activities in fern gametophytes directly contribute to this ecological differentiation and radiation in the future studies.

## Supplementary data

**Supplementary Figs. S1-S11**

Supplementary Fig. S1. Diagrams illustrating an apical initial and a lateral meristem in fern gametophytes.

Supplementary Fig. S2. Representative images of *P. vittata* gametophytes at 8 DAI and 10 DAI.

Supplementary Fig. S3. Computational segmentation and quantification of the confocal images.

Supplementary Fig. S4. Confocal imaging and computational analysis of the early gametophyte development in *P. vittata*.

Supplementary Fig. S5. The establishment of apical initials in *P. vittata* gametophytes.

Supplementary Fig. S6. The proliferation of *P. vittata* gametophytes.

Supplementary Fig. S7. The maintenance of apical initials in *P. vittata* gametophytes.

Supplementary Fig. S8. The initiation and proliferation of lateral meristems in *P. vittata* gametophytes.

Supplementary Fig. S9. The maintenance of apical initials and lateral meristems in *P. vittata* gametophytes.

Supplementary Fig. S10. Two representative *P. vittata* gametophytes at 26 DAI.

Supplementary Fig. S11. Computational segmentation and quantification of the confocal images.

**Supplementary Tables S1-S8**

Supplementary Table S1. Summary of different gametophytes of *P. vittata* imaged from 3 DAI to 32 DAI.

Supplementary Table S2. Area quantification of each segmented cell from the *P. vittata* gametophyte shown in Supplementary Fig. S3 and Fig. 2S.

Supplementary Table S3. Area quantification of each segmented cell from the *P. vittata* gametophyte shown in Supplementary Fig. S11 A and Fig. 2K.

Supplementary Table S4. Area quantification of each segmented cell from the *P. vittata* gametophyte shown in Supplementary Fig. S11 B and Fig. 2M.

Supplementary Table S5. Area quantification of each segmented cell from the *P. vittata* gametophyte shown in Supplementary Fig. S11 C and Fig. 2O.

Supplementary Table S6. Area quantification of each segmented cell from the *P. vittata* gametophyte shown in Supplementary Fig. S11 D and Fig. 2Q.

Supplementary Table S7. Area quantification of each segmented cell from the *P. vittata* gametophyte shown in Supplementary Fig. S11 E and Fig. 2U.

Supplementary Table S8. Area quantification of each segmented cell from the *P. vittata* gametophyte shown in Supplementary Fig. S11 F and Fig. 2W.

## Acknowledgements

The authors thank the Purdue Bindley Bioscience Facility for the access of the ZEISS LSM880 confocal microscope. The authors also thank Prof. Elliot Meyerowitz at Caltech for the support and encouragement. This work was supported by Purdue University start-up and funds from Purdue Center for Plant Biology (to YZ) and by the NSF-IOS 1931114 (to JB and YZ).

## Author contributions

YZ conceived the research direction, XW performed the experiments, XW, AY, JB SM, SZ, and YZ analyzed and discussed the experimental results, AY performed computational analysis and quantification, YZ supervised the experiments, XW, JB and YZ wrote the manuscript, AY, SM and SZ revised the manuscript, and all authors approved the manuscript. Authors declare no conflict of interest.

