## Supplementary file for "Timing of meristem initiation and maintenance determines the morphology of fern gametophytes"

Supplementary Table S8. Area quantification of each segmented cell from the *P. vittata* gametophyte shown in Supplementary Fig. S11 F and Fig. 2W.

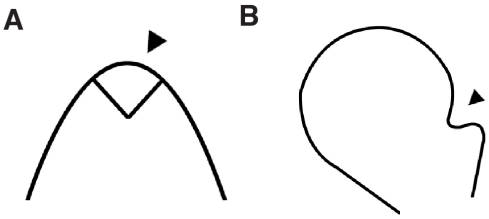


**Supplementary Fig. S1. (A-B) Diagrams illustrating an apical initial (A) and a lateral meristem (B) in fern gametophytes.**

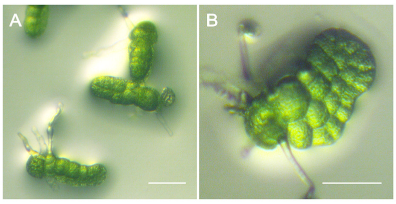


**Supplementary Fig. S2. (A-B) Representative images of *P. vittata* gametophytes at 8 (A) and 10 (B) DAI.** Scale bar: 10 μm (A, B).

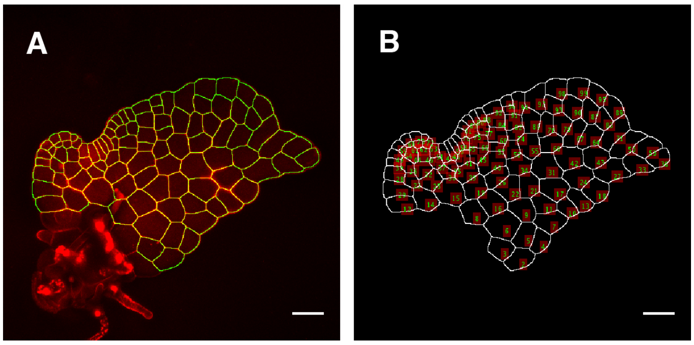


**Supplementary Fig. S3. Computational segmentation and quantification of the confocal images.** (A) The overlay of the confocal image (red) and the computational segmentation (yellow) of cell wall. (B) Each segmented cell with a unique ID in the gametophyte and the area of each segmented cell is automatically quantified. Scale bar: 50 μm. The image of one gametophyte (in Fig. 2 S-T) is shown here as an example. The quantified area of each cell (with the ID) from this sample is shown in Supplementary Table S2. All gametophyte images in Figs. 2, 4-7 and Supplementary Figs. S4-S9 were analyzed and quantified using the same procedure.

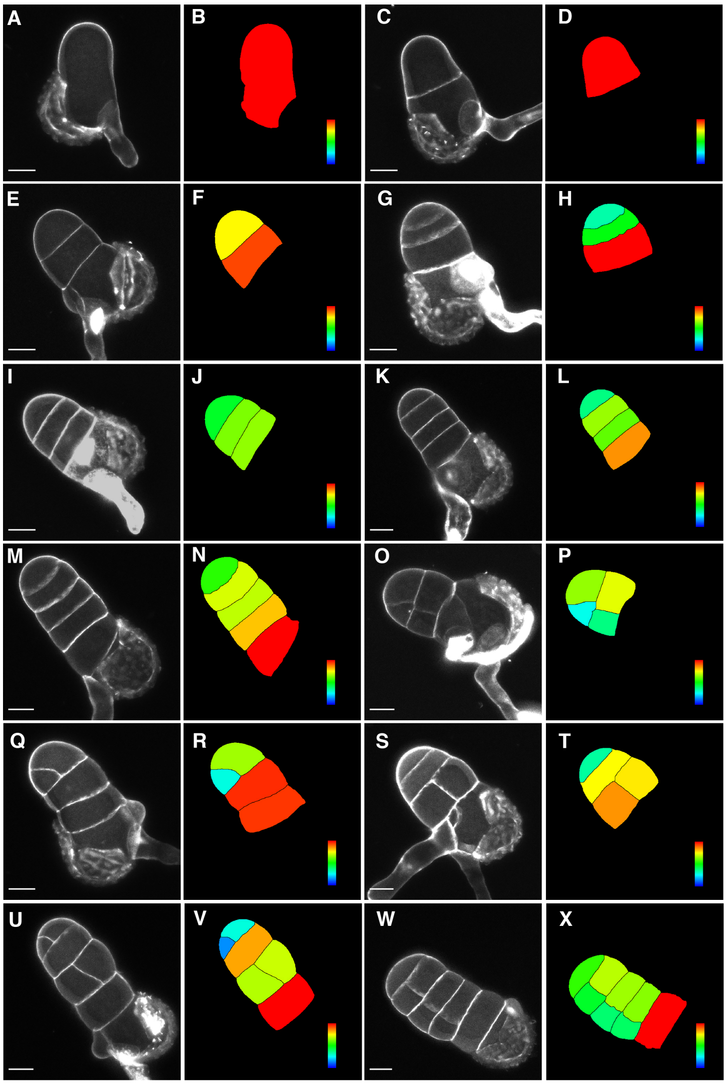


**Supplementary Fig. S4. Confocal imaging and computational analysis of the early gametophyte development in *P. vittata*.** Snapshots of different samples represented the spore germination before the establishment of an apical initial and the early gametophyte development. 12 gametophytes (A, C, E, G, I, K, M, O, Q, S, U, W) were stained and imaged through the laser scanning confocal microscopy. (B, D, F, H, J, L, N, P, R, T, V, X) show the computational segmentation and cell size quantification of confocal images in (A, C, E, G, I, K, M, O, Q, S, U, W). (A, C) show two gametophytes at 4 DAI, (E, G, I) show three gametophytes at 5 DAI, (K, M, Q, U) show four gametophytes at 6 DAI, (O, S) show two gametophytes at 7 DAI, and (W) shows a gametophyte at 8 DAI. Rhizoids was not analyzed in this study. The basal cells associated with the rhizoid and/or spore coat were not analyzed either. Scale bar: 20 µm. Gray (A, C, E, G, I, K, M, O, Q, S, U, W): propidium iodide (PI) stain. Color bar in (B, D, F, H, J, L, N, P, R, T, V, X) indicates the quantified area of each segmented cell with the scale from blue (0) to red (at or above 1000 µm$^2$).

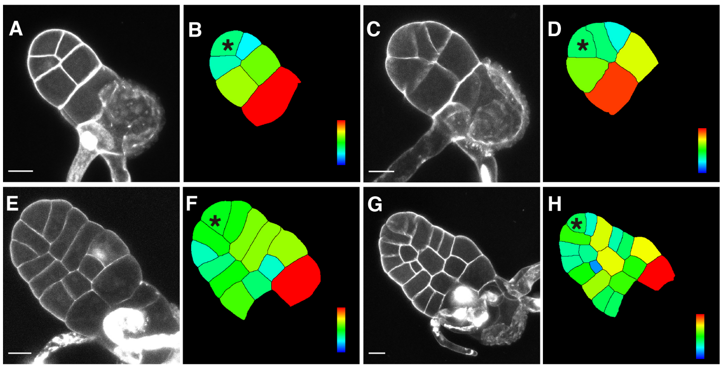


**Supplementary Fig. S5. The establishment of apical initials in *P. vittata* gametophytes.** Four gametophytes (A, C, E, G) were stained and imaged through the laser scanning confocal microscopy. (B, D, F, H) show the computational segmentation and cell size quantification of confocal images in (A, C, E, G), respectively. (A, C) show two gametophytes at 6 DAI, (E) shows a gametophyte at 9 DAI and (G) shows a gametophyte at 11 DAI. Stars indicate the apical initials. (A, C) show the earlies developmental stage when an apical initial can be clearly identified in the current growth condition. Scale bar: 20 µm. Gray (A, C, E, G): propidium iodide (PI) stain. Color bar in (B, D, F, H) indicates the quantified area of each segmented cell with the scale from blue (0) to red (at or above 1000 µm$^2$).

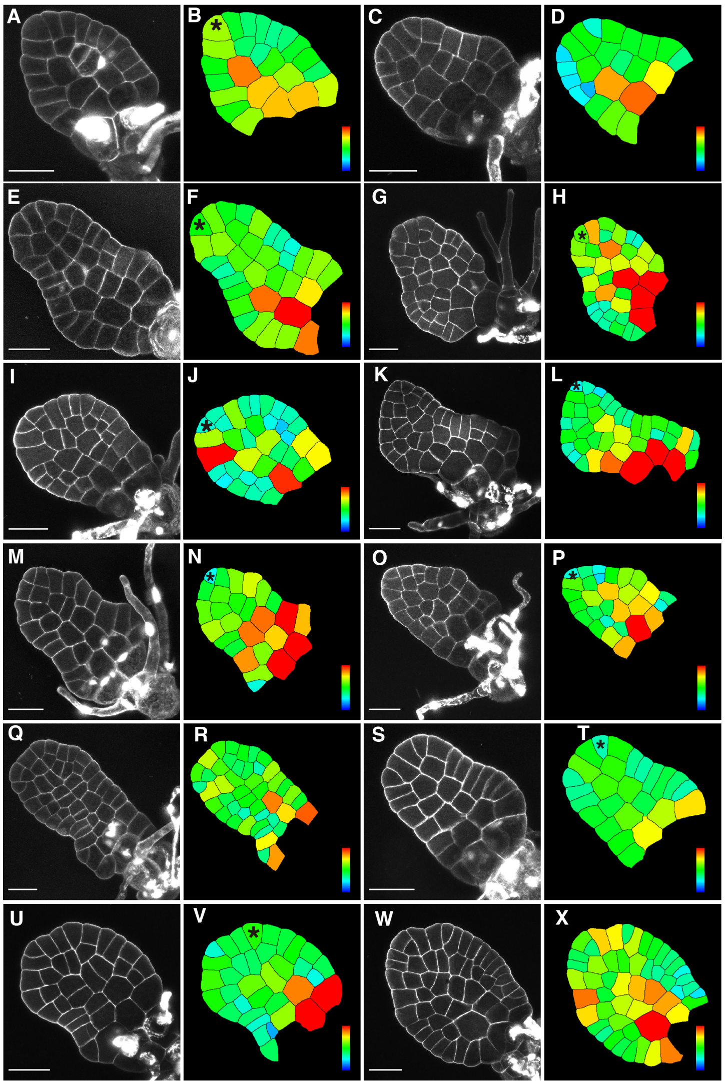


**Supplementary Fig. S6. The proliferation of *P. vittata* gametophytes.** 12 representative gametophytes (A, C, E, G, I, K, M, O, Q, S, U, W) were stained and imaged through the laser scanning confocal microscopy. (B, D, F, H, J, L, N, P, R, T, V, X) show the computational segmentation and cell size quantification of confocal images in (A, C, E, G, I, K, M, O, Q, S, U, W), respectively. (C) shows a gametophyte at 9 DAI, (A, E) show two gametophytes at 10 DAI, (S, U) show two gametophytes at 11 DAI, (I) shows a gametophyte at 12 DAI, (W) shows a gametophyte at 13 DAI, (G, M, O, Q) show a gametophytes at 14 DAI, and (K) shows a gametophyte at 15 DAI. Stars indicate the apical initials. Scale bar: 50 μm. Gray (A, C, E, G, I, K, M, O, Q, S, U, W): propidium iodide (PI) stain. Color bar in (B, D, F, H, J, L, N, P, R, T, V, X) indicates the quantified area of each segmented cell with the scale from blue (0) to red (at or above 1000 $\mu m^2$).

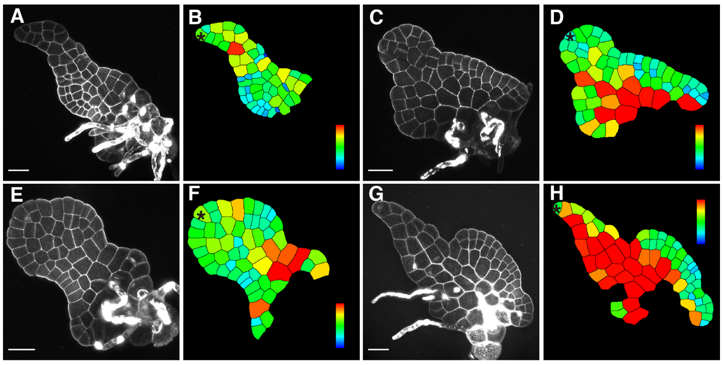



**Supplementary Fig. S7. The maintenance of apical initials in *P. vittata* gametophytes.** Four representative gametophytes maintaining the apical initials (A, C, E, G) were stained and imaged through the laser scanning confocal microscopy. (B, D, F, H) show the computational segmentation and cell size quantification of confocal images in (A, C, E, G), respectively. (A, C, E) show three gametophytes at 15 DAI, and (G) shows a gametophyte at 16 DAI. Stars indicate the apical initials. Scale bar: 50 µm. Gray (A, C, E, G): propidium iodide (PI) stain. Color bar in (B, D, F, H) indicates the quantified area of each segmented cell with the scale from blue (0) to red (at or above 1000 $\mu m^2$).

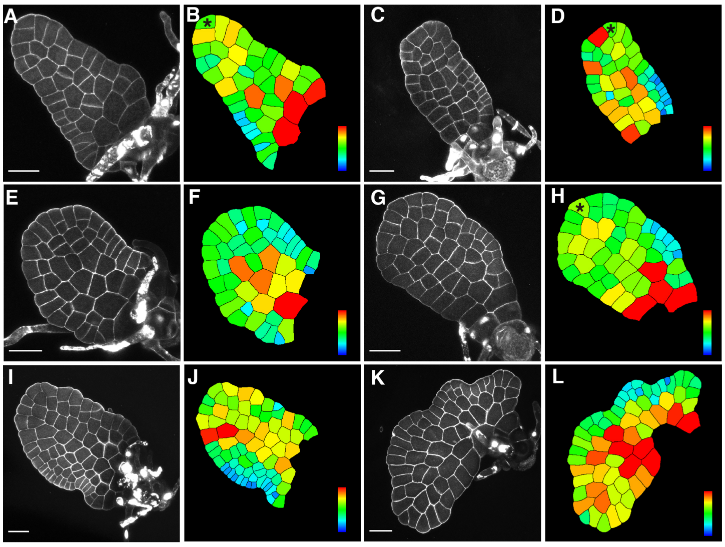



**Supplementary Fig. S8. The initiation and proliferation of lateral meristems in *P. vittata* gametophytes.** Six representative gametophytes (A, C, E, G, I, K) were stained and imaged through the laser scanning confocal microscopy. (B, D, F, H, J, L) show the computational segmentation and cell size quantification of confocal images in (A, C, E, G, I, K), respectively. Stars indicate the apical initials. (E) shows a gametophyte at 12 DAI, (A, G) show two gametophytes at 13 DAI, (C) shows a gametophyte at 15 DAI, (I) shows a gametophyte at 17 DAI, and (K) shows a gametophyte at 20 DAI. Scale bar: 50 μm. Gray (A, C, E, G, I, K): propidium iodide (PI) stain. Color bars indicate the quantified area of each segmented cell with the scale from blue (0) to red (at or above 1000 $\mu m^2$) in (B, D, F, H) and with the scale from blue (0) to red (at or above 1300 $\mu m^2$) in (J, L).

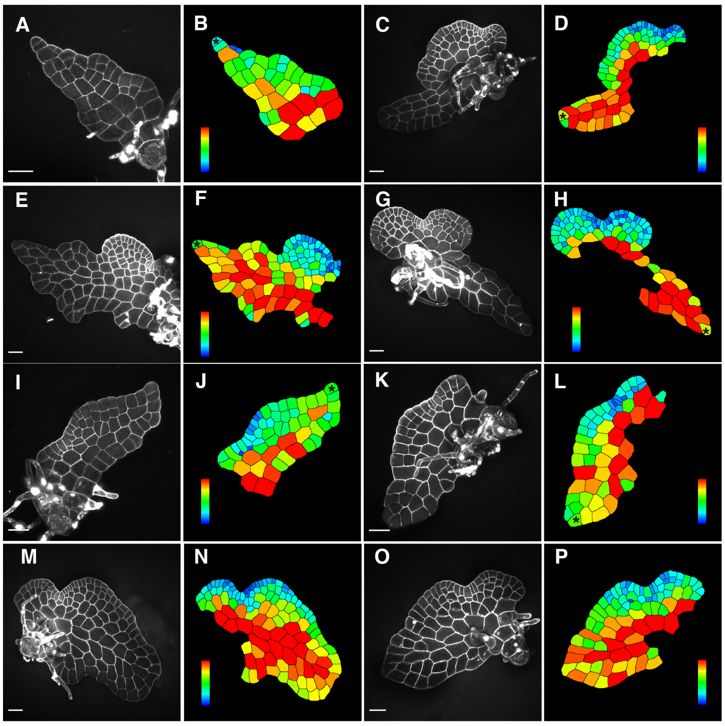


**Supplementary Fig. S9. The maintenance of apical initials and lateral meristems in *P. vittata* gametophytes.** The eight representative gametophytes (A, C, E, G, I, K, M, O) were stained and imaged through the laser scanning confocal microscopy. (B, D, F, H, J, L, N, P) show the computational segmentation and cell size quantification of confocal images in (A, C, E, G, I, K, M, O), respectively. Stars indicate the triangular shape apical initials. (A) shows a gametophyte at 16 DAI, (K) shows a gametophyte at 18 DAI, (I) shows a gametophyte at 19 DAI, (C) shows a gametophyte at 22 DAI, (E, G, M) show three gametophytes at 23 DAI, and (O) shows a gametophyte at 29 DAI. Scale bar: 50 μm. Gray (A, C, E, G, I, K, M, O): propidium iodide (PI) stain. Color bars indicate the quantified area of each segmented cell with the scale from blue (0) to red (at or above 1000 μm$^2$) in (B), with the scale from blue (0) to red (at or above 1300 μm$^2$) in (D, J, L), and with the scale from blue (0) to red (at or above 1600 μm$^2$) in (F, N, H, P).

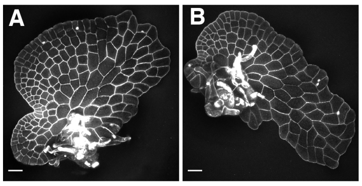


**Supplementary Fig. S10. Two representative *P. vittata* gametophytes at 26 DAI.** Two gametophytes were stained and imaged through the laser scanning confocal microscopy. (A) represents one type (Cordate, Ceratopteris like) and (B) represents the other type (Non-cordate, different from Ceratopteris) at 26 DAI, both of which are summarized in Table 1. Scale bar: 50 μm.

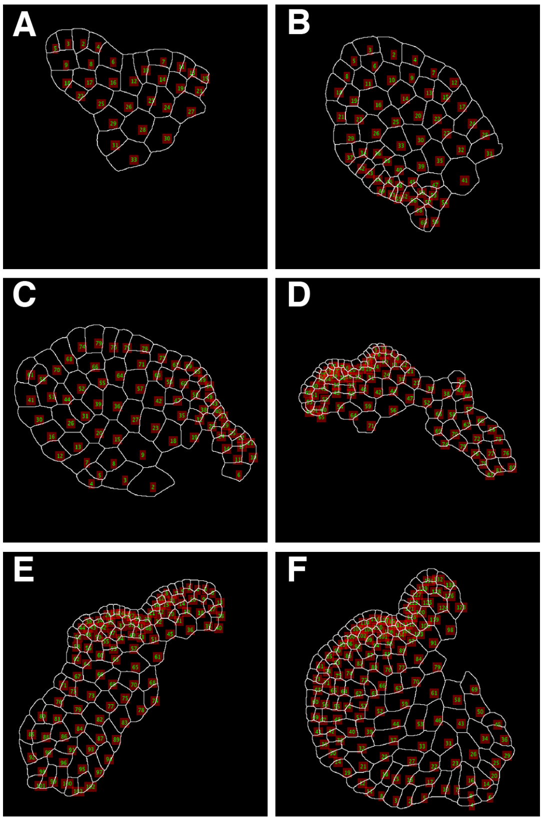

**Supplementary Fig. S11. Computational segmentation and quantification of the confocal images.** (A) The analyzed image of the gametophyte in Fig. 2K, including each segmented cell with a unique ID. The quantified area of each cell (with the ID) is shown in Supplementary Table S3. (B) The analyzed image of the gametophyte in Fig. 2M, including each segmented cell with a unique ID. The quantified area of each cell (with the ID) is shown in Supplementary Table S4. (C) The analyzed image of the gametophyte in Fig. 2O, including each segmented cell with a unique ID. The quantified area of each cell (with the ID) is shown in Supplementary Table S5. (D) The analyzed image of the gametophyte in Fig. 2Q, including each segmented cell with a unique ID. The quantified area of each cell (with the ID) is shown in Supplementary Table S6. (E) The analyzed image of the gametophyte in Fig. 2U, including each segmented cell with a unique ID. The quantified area of each cell (with the ID) is shown in Supplementary Table S7. (F) The analyzed image of the gametophyte in Fig. 2W, including each segmented cell with a unique ID. The quantified area of each cell (with the ID) is shown in Supplementary Table S8.
